# Fluorescent protein expression as a proxy of bacterial fitness in a high throughput assay

**DOI:** 10.1101/2020.12.01.399113

**Authors:** Rudolf O Schlechter, Evan J Kear, Daniela M Remus, Mitja NP Remus-Emsermann

## Abstract

Bacterial growth is classically assessed by measuring the increase in optical density of pure cultures in shaken liquid media. Measuring growth using optical density has severe limitations when studying multistrain interactions as it is not possible to measure the growth of individual strains within mixed cultures. Here we demonstrated that constitutively expressed fluorescent proteins can be used to track the growth of individual strains in different liquid media. Fluorescence measurements were highly correlated with optical density measurements and cell counts. This allowed us to assess bacterial growth not only in pure cultures, but also in mixed bacterial cultures and determine the impact of competitors on a focal strain, thereby assessing relative fitness. Furthermore, we were able to track the growth of two different strains simultaneously by using fluorescent proteins with differential excitation and emission wavelengths. Bacterial densities measured by fluorescence yielded more consistent data between technical replicates than optical density measurements. Our setup employs fluorescent microplate readers that allow for high throughput and replication.

**Importance:** We expand on an important limitation of the concept of measuring bacterial growth which is classically limited to one strain at a time. By adopting this approach, it is possible to measure growth of several bacterial strains simultaneously in high temporal resolution and in a high throughput manner. This is important to investigate bacterial interactions such as competition and facilitation.

## Introduction

Measurement of bacterial growth in liquid culture is a central paradigm for microbiology. Growth is specific for every bacterial strain and growth rates are determined as part of every characterisation of novel bacterial species or to describe the impact of mutations on bacterial fitness (1). Classically, growth is assessed by determining changes in turbidity and absorbance of liquid cultures using standards such as the McFarland standard (2) or by taking turbidity measurements in a spectrophotometer (3). These methods can be used to determine growth rates and predict cell numbers, maximal optical density (OD_600nm_) of individual strains, or whole communities in liquid medium. However, they suffer from limitations as spectrophotometers require bacterial cells to be well mixed without any formed clumps that would increase light scattering or cause heterogeneous turbidity (4). Thus, measuring growth of bacterial species that form aggregates in liquid media can be problematic. Furthermore, it is not possible to determine the growth of more than one bacterial strain in the same culture since only their combined optical density can be assessed. Alternatively to optical density, fluorescence dyes have been used to determine cell densities, e.g., the addition of the DNA intercalating dye acridine orange has been used to determine cell densities (5). This is possible by fluorometric analysis of liquid samples. Next to chemical dyes, fluorescent proteins can be detected in fluorometric analysis.

The green fluorescent protein (GFP) has been used in molecular microbiological experiments since the nineties (6, 7). Since then, many additional fluorescent proteins have been discovered or developed (e.g., (8, 9)). Many of these new proteins have dramatically different properties such as improved brightness and photostability, but also different spectral excitation and emission properties (10–12). Using GFP as a proof of concept, it has been demonstrated that constitutively expressed fluorescent proteins can be used to predict growth and colony forming units of *Pseudomonas aeruginosa* (13). This encouraged us to pursue a similar approach to predict the growth of bacteria in different liquid media and in co-culture with other differentially tagged bacteria.

Here, we demonstrated the use of different constitutively expressed fluorescence proteins as a means to measure bacterial growth. Furthermore, we used proteins with differential excitation and emission spectra (14) to allow the measurement of several populations at the same time. We performed our experiments in 96-well microtiter plates and microtiter plate readers, which allows for high replication and time series monitoring of fluorescent signals and cell growth in different growth conditions. By combining these measurements with a series of controls, we demonstrate that our system allows us to determine the strength of interaction and relative fitness of bacterial strains in competition.

## Results

### Growth of Pe::red in nutrient broth

To highlight the advantage of fluorescence intensity measurements over optical density measurements, we followed the growth of *Pantoea eucalypti* 299R labelled with a constitutively expressed red fluorescent protein (Pe::red) in five replicated cultures using optical density and fluorescence intensity in nutrient broth (NB) (Figure 1 A). The optical density increased exponentially for ∼5 h (Figure 1 B) to an optical density of 1.2. Afterwards, the culture entered a stationary phase for roughly 20 h followed by a rapid decline in optical density. The corresponding fluorescence increased exponentially for ∼10 h (Figure 1 B) and to a fluorescence intensity of 20,000 arbitrary units (a.u.). Afterwards, the rate with which fluorescence intensity increased was slightly reduced, indicating that cell growth decreased. After 20 h, the fluorescence intensity peaked at about 35,000 a.u. After a small reduction of fluorescence, suggesting cell lysis, the fluorescence increased again till the end of the experiment at 48 h. Growth curves obtained through optical density deviated even though the cultures were seeded from the same original culture and were diluted to the same initial optical density. The deviation was up to 0.25 OD_600nm_ which is more than 10% between the individual cultures. By contrast, the fluorescence of the individual cultures exhibited much lower deviations showing that measuring fluorescence yielded more consistent results.

**Figure 1.**
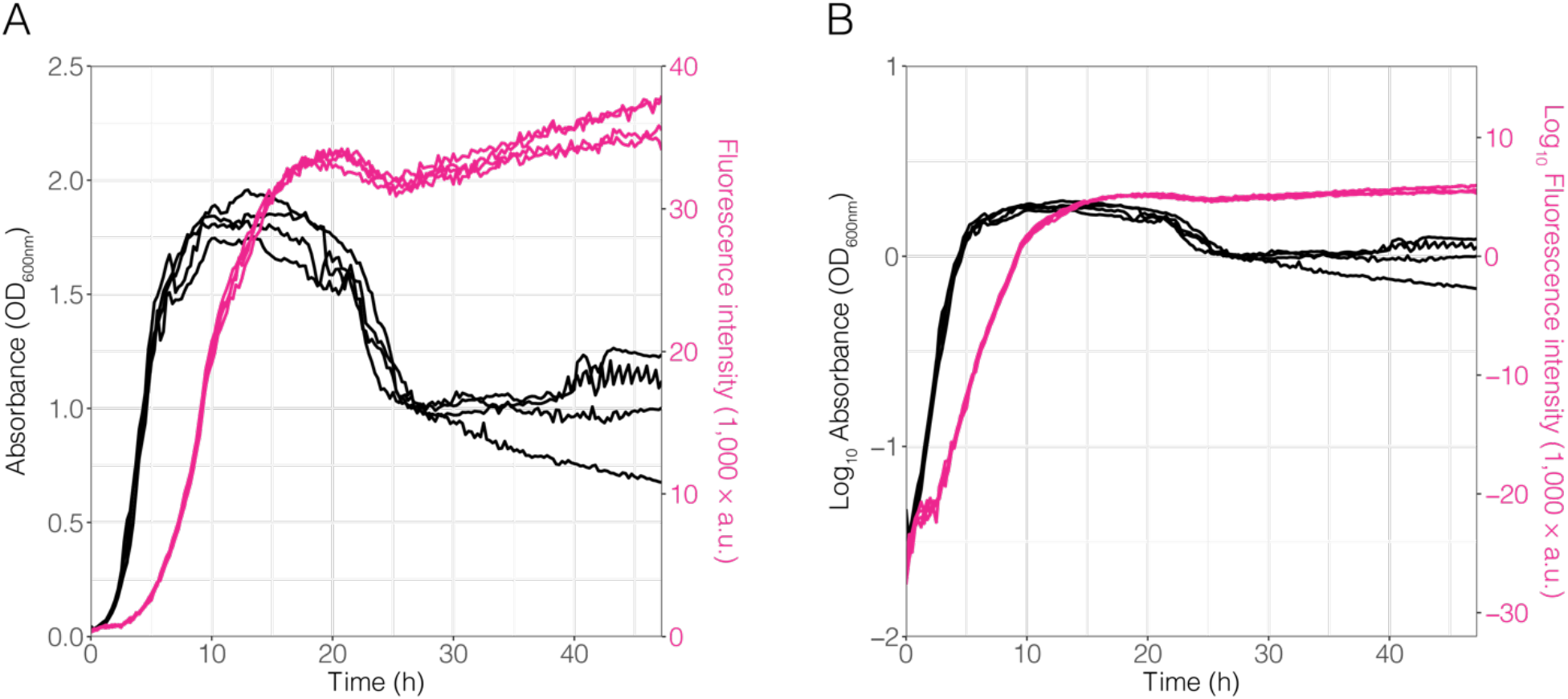
Pe::red growth in nutrient broth. Growth of Pe::red in nutrient broth in a 96-well microtiter plate was tracked using optical density (black) and fluorescence intensity (magenta). Optical density was measured as the absorbance of the culture at 600 nm (OD_600nm_) and red fluorescence was measured at 613–626 nm (a.u.: arbitrary units). Data was expressed in **(A)** linear and **(B)** logarithmic scale.

Furthermore, after 20 h of growth in 96-well plates, optical density of Pe::red decreased while red fluorescence steadily increased over time (Figure 1 A). To determine whether cell lysis or flock formation were associated with this growth pattern, independent cultures of Pe::red grown in NB were sampled over time and observed under the microscope for qualitative analysis. At early time points —that is, from 0 to 4 h—, most cells were planktonic and emitted fluorescence (Figure S1). However, we observed both cell lysis and flock formation at 24 h, as a proportion of cells showed morphological changes in the phase contrast, indicative of cell lysis, and loss of fluorescence (Figure S1), while other cell subpopulations aggregated (Figure S2).

### Growth of Pe::red in diluted NB

After inoculation into different concentrations of NB, Pe::red grew up to different final optical densities, while the initial lag phase and growth were similar (Fig. 2 A). As expected, the final optical density was directly dependent on the concentration of the NB medium. Depending on the available resources, the different cultures reached their stationary phases at different times and densities. Pe::red grown on 6.75% strength NB reached its stationary phase after ∼2.5 h while cultures growing on 12.5, 25 and 50% strength NB reached their stationary phase after 5 h. Cultures growing on higher concentrations of NB reached their stationary phase between 7.5 and 10 h. It is noteworthy to mention the large standard deviations and irregular curves, especially after the cultures entered their respective stationary phase. This was more pronounced at higher optical densities. After 15 h, some of the cultures exhibited a small decline in optical density.

**Figure 2.**
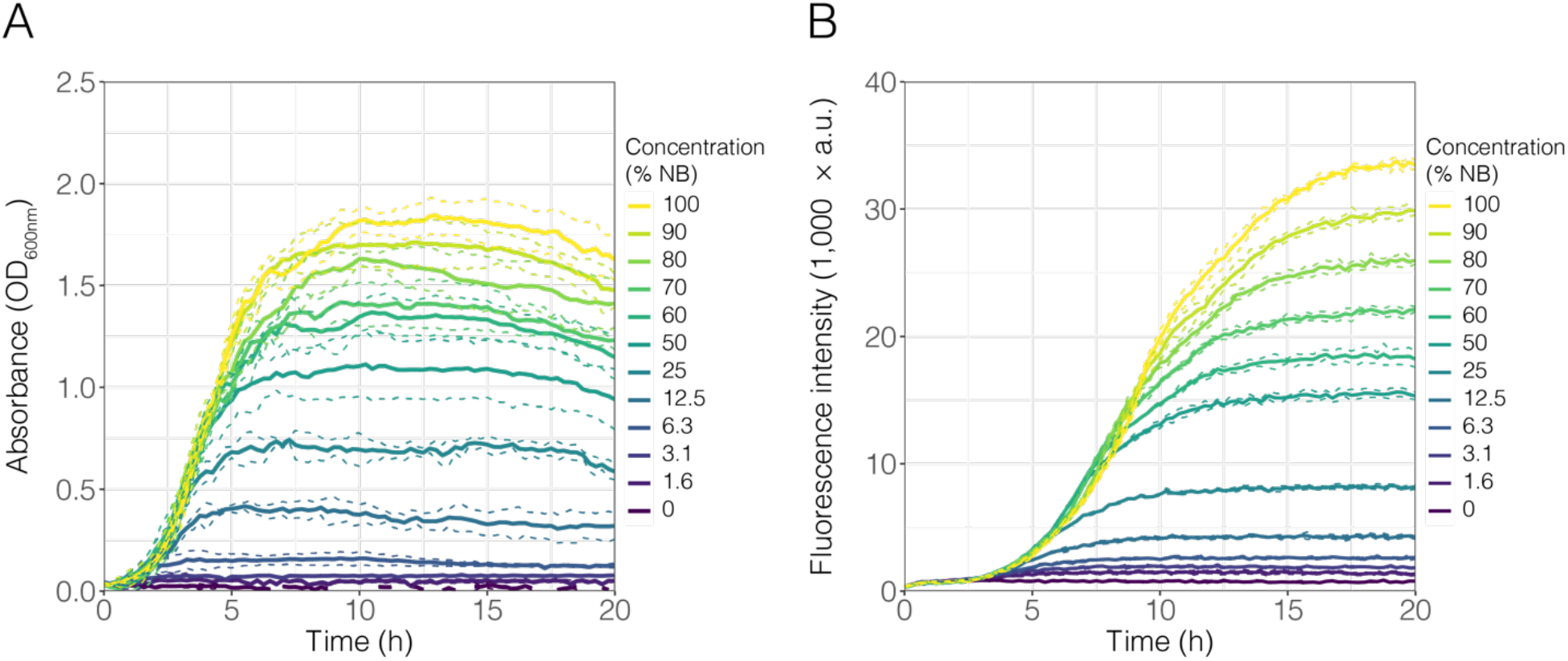
Growth of Pe::red on different concentrations of nutrient broth. Pe::red was inoculated into 96-well microtiter plates containing increasing concentrations of nutrient broth (NB), ranging from 0 to 100% v/v strength (see legend). Growth was measured by **(A)** optical density at 600 nm (OD_600nm_) and **(B)** fluorescence intensity emission (a.u.: arbitrary units) as a proxy. In both cases, continuous lines represent the mean of four biological replicates and dashed lines represent the standard deviation.

For fluorescence intensity measurements, the same cultures exhibited similar increases in fluorescence over time in increasing concentrations of NB as the measurements of optical densities (Fig. 2 B). As expected, the increase in fluorescence lagged the increase in optical density, likely due to the fluorescent protein maturation rate. The overall ranking of maximal fluorescence was the same as for the optical density measurements. The variability between the replicates was minimal in comparison to the optical density data. The fluorescence data did not exhibit a decline compared to the optical density data and remained stable till the end of the experiment after 20 h.

### Correlating bacterial growth with optical density and fluorescence intensity

Next, we tested if constitutively expressed fluorescence signals can be used as a proxy for growth. To that end, we used different measures of the optical density and fluorescence data extracted from Figure 2: the area under the curve (AUC), the maximal value, and the final value for each curve. We fitted these data into simple linear regression models of fluorescence and optical density to determine the parameters that gave the best fit by comparing the adjusted R^2^ and Pearson’s correlations (Table 1). Generally, we observed that every linear regression fitted the data with an R^2^ >0.95. The AUC of fluorescence data was the best predictor for every parameter of optical density used (i.e., AUC, maximum value, and final value), resulting in the best fits (R^2^ = 0.98) and highest Pearson’s correlation values (0.99) (Table 1).

**Table 1.**
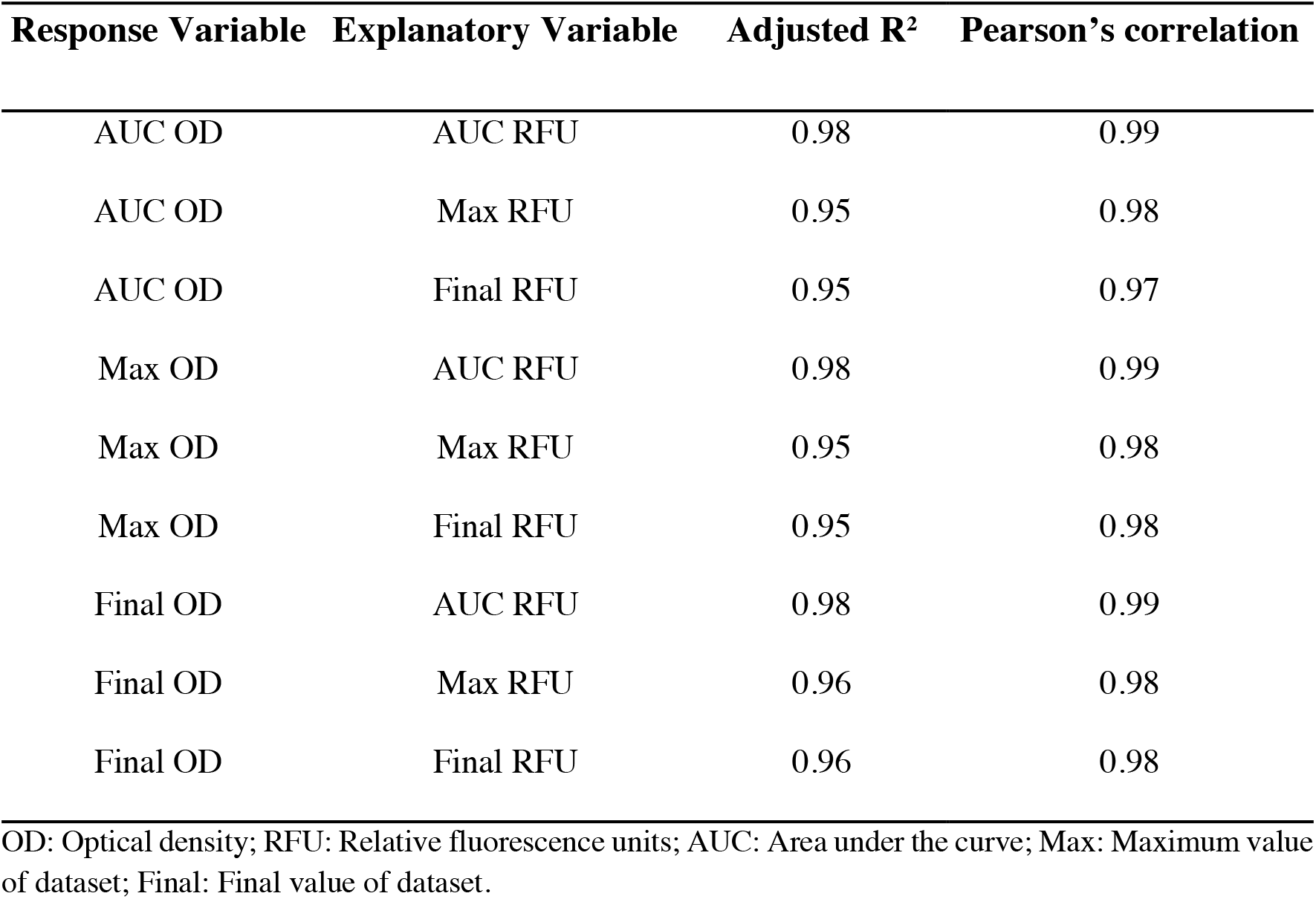
Parameters derived from linear regression models and correlation between the fluorescence and absorbance measurements presented in Figure 2.

In addition, we compared the optical density, fluorescence, and colony counts of two independent Pe::red NB cultures that were grown in Erlenmeyer flasks that were sampled over time. To test the change in fluorescence intensity in Pe::red over time, the fluorescence emission of individual Pe::red cells was measured and analysed at different time points. We observed that from 0 to 4 h, single cells exhibited a statistically significant, albeit small, increase of fluorescence (Figure S3). However, the low coefficient estimates derived from generalised linear models for each replicate together with the low goodness-of-fit for each model (Replicate 1: intercept = 940.85 a.u.; coefficient estimate = 0.18, adjusted R^2^ = 0.0079; Replicate 2: intercept = 532.75; coefficient estimate = 3.25, adjusted R^2^ = 0.13) suggested that growth time had only a small effect on single-cell fluorescence intensity, where about 10% of the data can be explained by these models. The results indicate that the fluorescence of individual cells increased at a very small rate at early growth phases and that the increase in fluorescence in a population can be associated with an increase in the population density instead of an increase in protein maturation rate. A strong correlation between fluorescence and optical density was observed between Pe::red cultures growing in NB sampled at different time points (Table 2, Figure 3 A). Similarly, both optical density and fluorescence intensity also correlated with bacterial counts, measured as CFU mL^-1^ (Figure 3 B and 3 C, respectively).

**Table 2.** Parameters derived from linear regression models and correlation between the fluorescence and absorbance measurements presented in Figure 3.

| Response Variable | Explanatory Variable | Adjusted R<sup>2</sup> | Pearson's correlation |
| --- | --- | --- | --- |
| Fluorescence intensity | Optical density | 0.84 | 0.92 |
| Bacterial counts* | Fluorescence intensity* | 0.78 | 0.89 |
| Bacterial counts* | Optical density* | 0.84 | 0.93 |

\* Data was transformed to logarithmic scale (log<sub>10</sub>).

**Figure 3.**
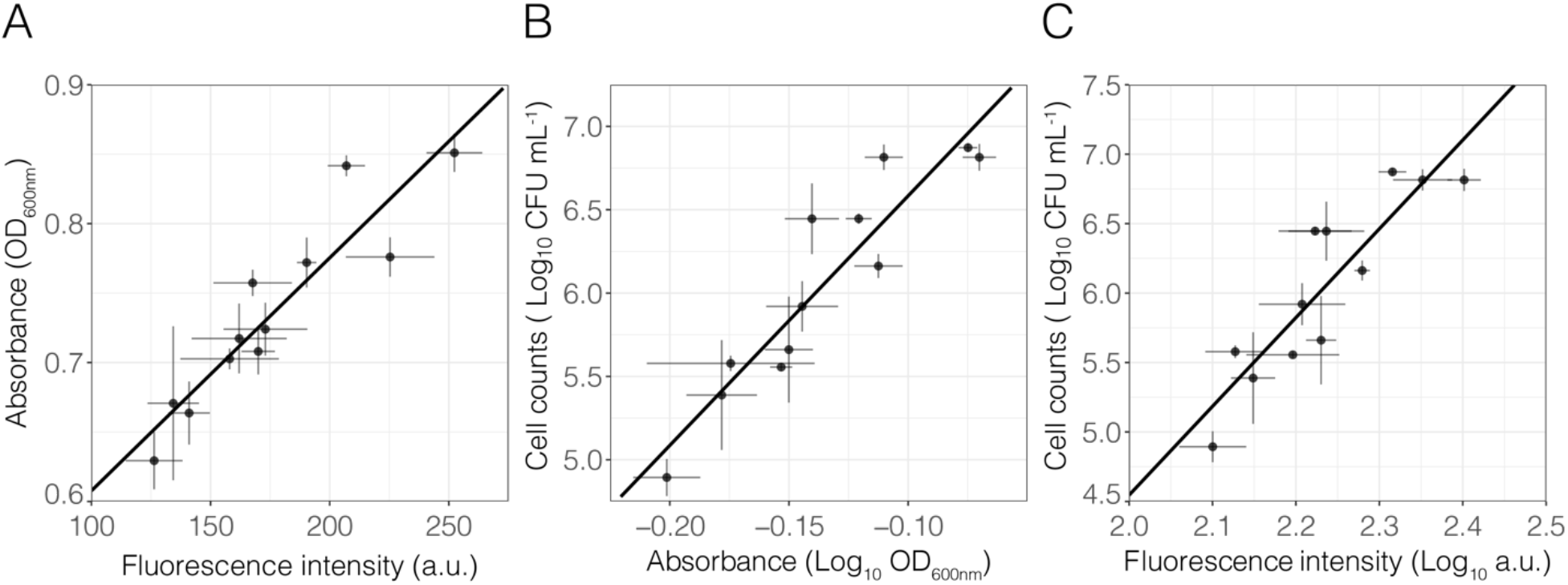
Relationships between fluorescence, absorbance and cell counts. Fluorescently tagged Pe::red was grown in shaken conical flasks. Samples were taken after 0, 0.75, 1.50, 2.50, 3.25, and 4 h post-inoculation. Each point represents the average of three technical replicates for each timepoint and each biological replicate (n = 2), with error bars representing SD. Lines depict simple linear regression models. **(A)** Fluorescence *vs* absorbance (linear regression, *p*-value < 0.05, R_2_ = 0.84, Pearson’s correlation = 0.92), **(B)** Absorbance *vs* cell count (linear regression, *p*-value < 0.05, R_2_ = 0.84, Pearson’s correlation = 0.93), **(C)** Fluorescence *vs* cell count (linear regression, *p*-value < 0.05, R_2_ = 0.78, Pearson’s correlation = 0.89). In **(B)** and **(C)**, cell counts were determined as CFU mL_-1_ and data is represented in logarithmic scale (log_10_). A.u. = arbitrary units, OD_600nm_ = optical density at 600 nm.

Finally, we evaluated the correlation between fluorescence and optical density in strains other than Pe::red and in different growth media. To that end, fluorescently tagged strains, listed in Table 3, were incubated in minimal medium (MM) supplemented with 31 different carbon sources, and both fluorescence emission and final OD_600nm_ were measured. We selected a wide range of phylogenetically different strains (from *Proteobacteria* and *Actinobacteria*) that constitutively express a red or green fluorescent protein. In general, we observed a statistically significant and positive correlation between the AUC of the fluorescence intensity curve of bacterial strains and the final OD_600nm_ of each liquid culture (Figure S4, GLMM, *p*-value <0.05, *pseudo*-R^2^ = 0.80, Pearson’s correlation = 0.72).

**Table 3.**
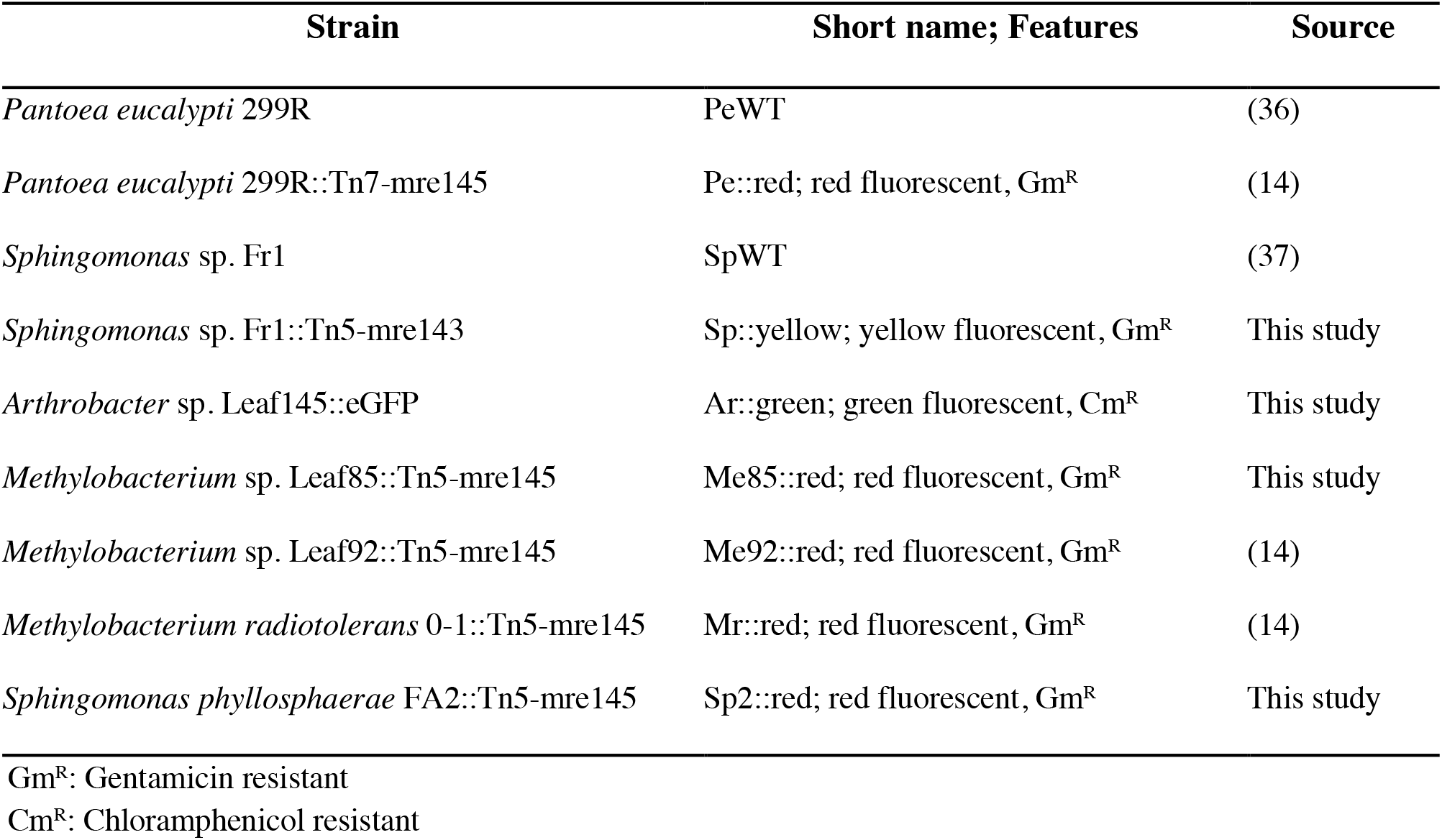
Bacterial strains used in this work

| Strain | Short name; Features | Source |
| --- | --- | --- |
| *Pantoea eucalypti* 299R | PeWT | (36) |
| *Pantoea eucalypti* 299R::Tn7-mre145 | Pe::red; red fluorescent, Gm<sup>R</sup> | (14) |
| *Sphingomonas* sp. Fr1 | SpWT | (37) |
| *Sphingomonas* sp. Fr1::Tn5-mre143 | Sp::yellow; yellow fluorescent, Gm<sup>R</sup> | This study |
| *Arthrobacter* sp. Leaf145::eGFP | Ar::green; green fluorescent, Cm<sup>R</sup> | This study |
| *Methylobacterium* sp. Leaf85::Tn5-mre145 | Me85::red; red fluorescent, Gm<sup>R</sup> | This study |
| *Methylobacterium* sp. Leaf92::Tn5-mre145 | Me92::red; red fluorescent, Gm<sup>R</sup> | (14) |
| *Methylobacterium radiotolerans* 0-1::Tn5-mre145 | Mr::red; red fluorescent, Gm<sup>R</sup> | (14) |
| *Sphingomonas phyllosphaerae* FA2::Tn5-mre145 | Sp2::red; red fluorescent, Gm<sup>R</sup> | This study |

Gm<sup>R</sup>: Gentamicin resistant Cm<sup>R</sup>: Chloramphenicol resistant

Consequently, data derived from the fluorescence signals allowed the tracking of growth in liquid cultures, showing that fluorescence can be used as an alternative to OD_600nm_.

### Growth of Pe::red and Sp::yellow in presence of different competitors

To determine the impact of different competitors on the growth of Pe::red and Sp::yellow, we inoculated mixed cultures into 96-well plates and covered the plates with a gas permeable foil to avoid evaporation during long-term incubation. Each fluorescently tagged strain was mixed with their respective wild type conspecific strain (Pe::red *vs*. PeWT and Sp::yellow *vs*. SpWT), and also a mix combining both strains (Pe::red *vs*. Sp::yellow), allowing us to track growth of each strain in parallel (Fig. 4 A). As control, fluorescent strains were grown in monoculture. Their respective fluorescence over time was determined every 10 minutes. As expected when grown without a competitor, Pe::red and Sp::yellow both reached the highest fluorescence intensity and hence, the highest cell density (Fig. 4 A, entire lines). When grown against their near-isogenic wild types, the fluorescence did not reach the same fluorescence levels depicting the impact of the competitor on the focal strain. We used the AUC of the growth curves as a proxy for bacteria abundance and to determine bacterial population sizes relative to their respective fluorescent monoculture. The relative AUC of the competition Pe::red *vs*. PeWT was on average 0.35 when normalised by the AUC of the Pe::red monoculture (Fig. 4 B). Since conspecifics were expected to show no fitness differences when competing against each other, we evaluated the fitness of independent Tn*7*-insertion Pe::red mutant strains. Results from this experiment showed that the relative fitness of Pe::red in competition with PeWT were consistent among strains and were usually 0.5 (Figure S5). The relative AUC of Sp::yellow *vs*. SpWT normalised by the AUC of the Sp::yellow monoculture was 0.49 and hence, Sp::yellow grew to almost exactly half the population density of the monoculture (Fig. 4 C). When competing Pe::red and Sp::yellow, the relative AUC of Pe::red and Sp::yellow was 0.79 and 0.22, respectively, when normalised by their respective monocultures (Fig. 4 B and C). Hence, Pe::red had a competitive advantage over Sp::yellow and the Sp::yellow was strongly impacted by the presence of Pe::red. Additionally, this competition assay was evaluated in MM supplemented with 0.2% w/v succinate, a resource that both strains can grow on. We observed a different pattern in which Pe::red in competition with Sp::yellow did not differ with Pe::red grown as monoculture (Figure S6 A), while Sp::yellow was affected by the presence of Pe::red (Figure S6 B). These results highlighted the applicability of this approach to different experimental setups to study the dynamics of bacterial interactions simultaneously and to address hypotheses regarding species interactions in defined growth conditions.

**Figure 4.**
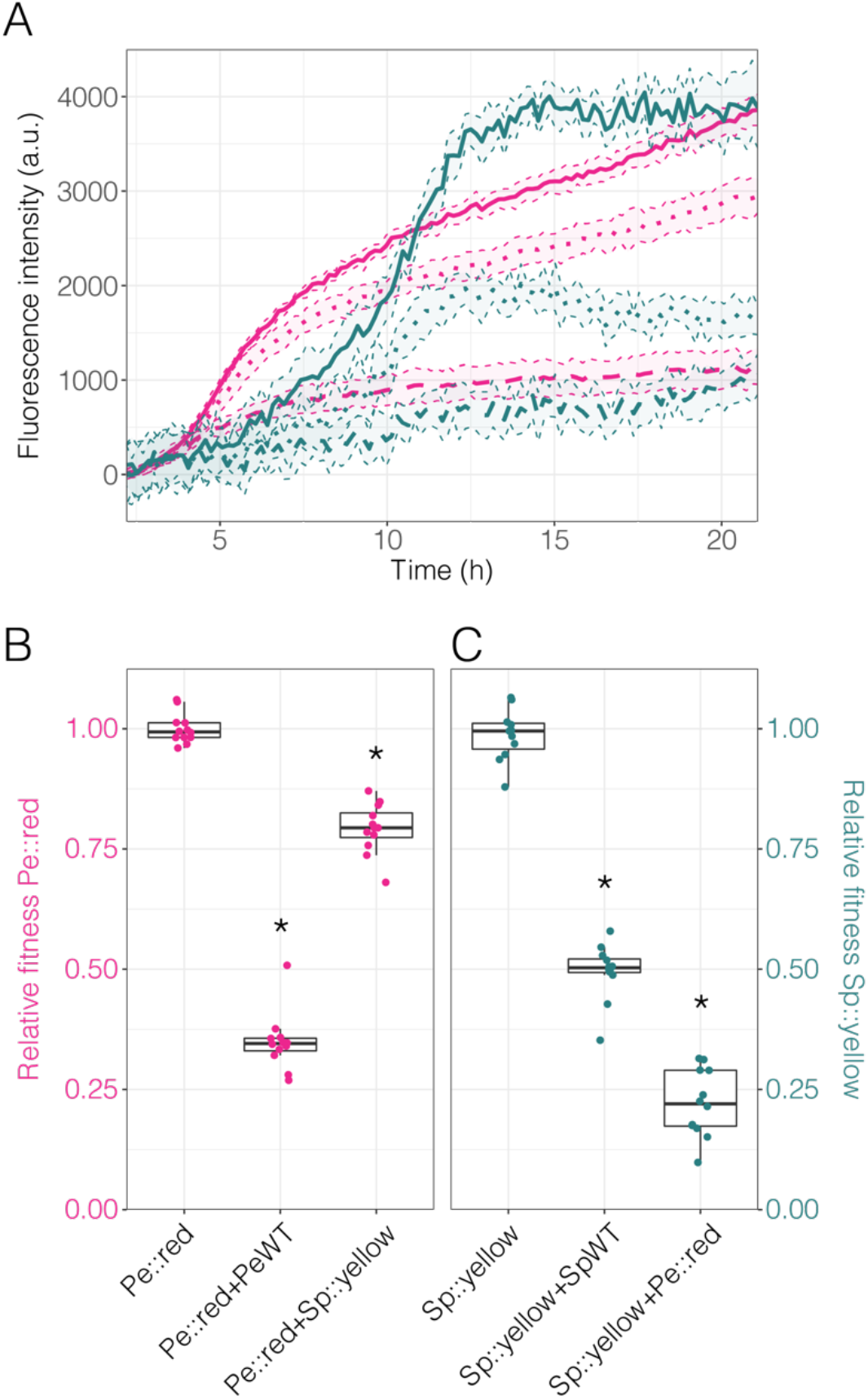
Competition assay in rich media. The fluorescently tagged bacterial strains Pe::red and Sp::yellow were competed in nutrient broth and their growth was estimated using fluorescence intensity. The data was background subtracted to normalise against the autofluorescence of the media. **(A)** Fluorescent intensity over time for each strain under different conditions. Magenta: red fluorescence emitted by Pe::red; dark green: yellow fluorescence emitted by Sp::yellow. Continuous lines represent monocultures of Pe::red or Sp::yellow. Broken lines represent each culture of Pe::red or Sp::yellow in competition with the respective wild type PeWT or SpWT strains. Dotted lines represent each culture of Pe::red and Sp::yellow in co-culture. Thin broken lines represent the standard deviation from the mean. **(B)** Relative fitness of Pe::red in monoculture, co-culture with PeWT, and co-culture with Sp::yellow. Fitness determined as the AUC derived from red fluorescence relative to Pe::red in monoculture. **(C)** Relative fitness of Sp::yellow in monoculture, co-culture with SpWT, and co-culture with Pe::red. Relative fitness of Sp::yellow was determined as the AUC derived from yellow fluorescence relative to Sp::yellow in monoculture. In **(B)** and **(C)**, groups were compared using one-way ANOVA test, where * = *p*-value < 0.05 compared to the monoculture (Bonferroni *post-hoc* test).

## Discussion

We have designed an experimental platform that allows tracking of bacterial growth in 96-well plates using fluorescence measurement as an alternative to optical density. This allowed us to track distinct populations in parallel and we provided an example of a mixed culture of two populations. Fluorescence measurements in plate readers also have the advantage of being less sensitive to cell aggregation than optical density, as several measurements of the same culture are taken and averaged, thereby integrating large areas of the sample and accounting for aggregates and less dense areas of the sample. In conjunction with controls, this approach allows for the measurement of competition between strains in a high throughput manner. With optimised detector systems such as filters or monochromators and well-chosen fluorescent proteins, it would be possible to track at least three differently tagged populations without complex experimental controls (e.g., cyan, yellow, and red fluorescent proteins).

We initially performed our experiments with *Pantoea eucalypti*, a fast-growing strain that does not form biofilms in shaken liquid culture. This allowed us to compare optical density and fluorescence intensity under defined conditions. Growth curves of optical density and fluorescence of bacterial cultures do not resemble each other perfectly, which is not surprising since the rate and maturation of fluorescent proteins is a combination of time consuming steps from DNA transcription, mRNA translation and protein maturation before the fluorescence signal can be detected (15). This process is however reproducible under controlled growth conditions. Furthermore, others have shown that by using a standard curve, it is possible to predict colony-forming units from fluorescence data (13). Our experiments have shown that the constitutive expression of fluorescent proteins is suitable to determine bacterial growth in shaking liquid cultures. Every measure that we tested correlated well with bacterial growth as measured by optical density and cell counts (Table 1 and 2).

Since growth assessed using fluorescence yields more consistent data than optical density in 96-well plates, it may also be useful in applications such as normalisation of fluorescence intensity of bacterial bioreporters (16). Using our workflow, it is possible to conveniently determine relative fitness of bacteria. Previously, the relative fitness of bacteria has been assessed using shared media reservoirs and filters to separate bacterial populations, which allowed for low throughput investigations of bacterial densities after the competition measured by optical density (17) or by determining the colony-forming units at different sampling times in shared media (e.g. (18, 19)). Both methods have in common that they cannot follow population dynamics at high resolution, such as early or late success or bacterial growth rate in presence of a competing species. By using continuous tracking of bacterial cultures in combination with control experiments that allow for normalisation we can infer changes to growth during all phases of an experiment.

To overcome evaporation of medium, which is one of the major drawbacks of 96-well plates that leads to strong edge effects and impacts optical density, we have used hydrophobic, gas-permeable foils to seal 96-well plates. The foils reduced evaporation to negligible amounts over the course of our experiments (up to two weeks, data not shown). At the same time, it also leads to reduced oxygen availability for bacterial growth and limits the ability to measure optical density due to turbidity of the foil and condensation of droplets on the foil. However, since the reduced oxygen availability impacts on the whole plate equally, competition treatments and respective control will be both similarly impacted, and the resulting relative fitness can still be assessed.

The experimental system is not without flaws and even though many bacterial strains exhibit negligible autofluorescence in liquid media, some exhibit strong autofluorescence, such as pseudomonads in minimal media (20). To be able to accurately assess the growth of strongly autofluorescent bacteria, it is necessary to determine the autofluorescence of the respective parental strain without the fluorescent protein expressed. One clear disadvantage of fluorescent proteins is that many bacterial strains are not amenable to genetic modification. However, several recent publications were able to increase the breadths of bacterial recipients that can successfully be manipulated using stable plasmids that do not require antibiotic pressure for medium term maintenance, or (transposon mediated) chromosomal integration (14, 21, 22). Our system should still be interesting for those hard to modify bacteria by competing them against known, fluorescently tagged organisms by determining the impact of the non-tagged organism on the tagged one.

## Conclusion

We have designed a convenient experimental platform that tracks bacterial growth by employing constitutively expressed fluorescent proteins. Our platform allows us to track several fluorescence signals simultaneously in a time-resolved manner and in high replications. It yields more robust results than classical turbidity measurements and can be used to determine competition in controlled conditions.

## Materials and Methods

### Strains and growth conditions

Strains used in this study are listed in Table 3. Bacteria were routinely grown on nutrient broth (NB, HiMedia) or nutrient agar (NA, HiMedia) supplemented with 15 mg L^-1^ gentamicin, 15 mg L^-1^ tetracycline, or 10 mg L^-1^ chloramphenicol where appropriate. Broth cultures were incubated at 30°C in a rotary shaker at 200 rpm. Agar plates were incubated at 30°C. Plasmid pMRE-Tn5-143 was used to transform *Sphingomonas* sp. Fr1 by conjugation, as described previously (14, 23), to constitutively express sYFP2 (24). Using the same method, plasmid pMRE-Tn5-145 was used to deliver constitutively-expressed fluorescent protein carrying transposons to *Sphingomonas* sp. Fr1, *Sphingomonas phyllosphaerae* FA2, and *Methylobacterium* sp. Leaf85 to constitutively express mScarlet-I (25). *Arthrobacter* sp. Leaf145 was transformed by sono-electroporation (26) using the plasmid pKGT-GFP, kindly gifted by Christine Smart (27), to constitutively express eGFP (28). *Methylobacterium* sp. Leaf92::Tn5-mre145, *Methylobacterium radiotolerans* 0-1::Tn5-mre145, and *Pantoea eucalypti* 299R::Tn7-mre145 were constructed elsewhere (14).

Minimal medium (MM; 1.62 g L^-1^ NH_4_Cl, 0.2 g L^-1^ MgSO_4_, 1.59 g L^-1^ K_2_HPO_4_, 1.8 g L^-1^ NaH_2_PO_4_ •2H_2_O, 15 g L^-1^ agar, with the following trace elements: 15 mg L^-1^ Na_2_EDTA_2_•H_2_O, 4.5 mg L^-1^ ZnSO_4_•7H_2_O, 3 mg L^-1^ CoCl_2_•6H_2_O, 0.6 mg L^-1^ MnCl_2_, 1 mg L^-1^ H_3_BO_3_, 3.0 mg L^-1^ CaCl_2_, 0.4 mg L^-1^ Na_2_MoO_4_ • 2H_2_O, 3 mg L^-1^ FeSO_4_ • 7H_2_O, and 0.3 mg L^-1^ CuSO_4_ • 5H_2_O (29)) was supplemented with 0.2% w/v of a carbon source. Carbon sources used were Sucrose, D-Glucose, D-Fructose, Galactose, D-Ribose, Xylose, L-Arabinose, D-Mannitol, Sorbitol, Glycerol, Pyruvate, L-Malate, Citrate, 2-Oxoglutarate, Succinate, Maleate, Fumarate, L-Threonine, L-Glutamine, L-Glutamate, L-Serine, L-Aspartate, L-Proline, L-Lysine, L-Isoleucine, L-Alanine, L-Valine, L-Tryptophan, L-Asparagine, GABA, and Methanol.

### Plate reader experiments

High-throughput growth experiments were carried out in 96-well microtiter plates (Costar). The 96-well plates were sealed with either a lid or with the hydrophobic gas permeable membrane 4ti-0516/96 (Brooks Life Sciences, Wotton, UK; gas permeability 0.6 m^-3^ m^-2^ day^-1^ and a water loss of 1 g m^-2^ day^-1^). For every experiment in microtiter plates, and unless stated otherwise, overnight bacterial cultures were used to seed 96-well microtiter plates to reach a volume of 200 µL per well. To that end, cultures were harvested by centrifugation at 6,000 × *g* and washed twice in 1 × phosphate buffered saline (PBS; 8 g L^-1^ NaCl, 0.24 g L^-1^ KCl, 1.42 g L^-1^ Na_2_HPO_4_, 0.24 g L^-1^ KH_2_PO_4_). Finally, the washed cultures were resuspended and diluted to a defined adjusted optical density at 600 nm (OD_600nm_). Plates were incubated in a FLUOstar Omega plate reader (BMG Labtech) at 30°C for up to five days, depending on the growth rate of each strain and growth media. The plates were shaken in “meander corner well shaking” mode at 300 rpm between each read. Readings of OD_600nm_ and fluorescence were measured in bottom optic mode. Fluorescence was measured in a 2-mm diameter circle and the average of eight measurements per well was recorded. Optical density and the individual fluorescence spectra were measured sequentially to minimise crosstalk between the fluorescent proteins. mScarlet fluorescence was excited at 567-587 nm and emission was measured at 613-626 nm. sYFP2 and eGFP fluorescence were excited at 475-492 nm and emissions were measured at 511-550 nm.

### Growth experiments

To establish a relationship between fluorescence and optical density, Pe::red was grown in different concentrations of NB in 96-well plates. To that end, washed culture of Pe::red was resuspended and diluted to an OD_600nm_ of 0.05. To each well, aliquots of 20 µL resuspended culture were added to 180 µL NB or NB diluted to 90, 80, 70, 60, 50, 25, 12.5, 6.75, 3.4, 1.7 or 0% strength, respectively. The plate was closed with a lid. Optical density at 600 nm and red fluorescence were measured every 15 minutes as described above.

Additionally, cultures of fluorescently tagged strains listed in Table 1 were used to seed microtiter plates containing MM supplemented with 0.2 % w/v of individual carbon sources. To that end, every washed culture was resuspended, diluted to an OD_600nm_ of 0.05, and 20 µL resuspended cultures were seeded into each well in triplicates for each growth medium. Red or green fluorescence was measured every 15 min and OD_600nm_ was measured at the end of the experiment (Final OD_600nm_).

### Competition experiments and fitness assessment

Overnight cultures were resuspended to an OD_600nm_ of 0.37. Four µL of the diluted cultures per well were inoculated into 192 µL NB or MM supplemented with 0.2% w/v succinate in triplicates to perform the competition experiments. Then, the following bacterial mixes were prepared: Pe::red + 4 µL medium (Pe::red monoculture), Pe::red + PeWT (near-isogenic Pe co-culture), Sp::yellow + 4 µL medium (Sp::yellow monoculture), Sp::yellow + SpWT (near-isogenic Sp co-culture), Pe::red + Sp::yellow. In all cases, 96-well plates were sealed with a hydrophobic gas permeable membrane, and OD_600nm_, red, and yellow fluorescence were measured every 15 minutes. A similar setup was used to identify potential fitness costs of Tn*7* insertions in Pe::red. To that end, four independent Tn*7* mutant strains were used to compete against a near-isogenic strain (PeWT) or Sp::yellow.

### Conical flasks experiment

Two independent Pe::red cultures were grown overnight in NB with gentamicin from single colonies. The overnight cultures were inoculated into 250 mL conical flasks containing NB and gentamicin, resulting in 80 mL of culture at an initial density of OD_600nm_ = 0.05. At 0, 0.75, 1.50, 2.50, 3.25, 4, and 24 h post-inoculation (hpi), samples of each culture were taken for optical density, red fluorescence, colony counts, and microscopy.

Firstly, 200 µL of culture were transferred into a 96-well plate containing the appropriate blanks. Endpoint measurements were taken for optical density and red fluorescence in triplicates.

Secondly, ten-fold serial dilutions with dilution factors ranging from 10^2^ to 10^7^ were prepared in triplicate, and 5 µL of each dilution was plated on NB containing gentamicin. Plates were incubated at 30 °C overnight and counted under a stereo microscope.

Lastly, an aliquot of each culture was fixed for microscopy. To that end, 1 mL of each of the two cultures was harvested at 15,000 × *g*, washed in 1 mL PBS, and resuspended in 50 µL 1 × PBS. Then, 150 µL paraformaldehyde solution (PFA; 4% w/v paraformaldehyde) was added. After 30 minutes, cells were harvested by centrifugation and resuspended in 50 µL 1 × PBS, to which 50 µL 100% v/v ethanol was added after the cells were resuspended. Fixed samples were stored at - 20 °C until processed.

### Microscopy

Slides were prepared by applying 4 µL of each fixed sample onto 0.1% w/v gelatin-coated microscope slides. Samples were air dried and mounted using 60% glycerol.

Fluorescent microscopy was carried out using a Zeiss AxioImager.M1 fluorescent widefield microscope at 1000 × magnification and phase contrast (EC Plan-Neofluar 100×/1.30 Ph3 Oil M27 objective) equipped with Zeiss filter set 43HE (BP 550/25-FT 570-BP 605/70), an Axiocam 506, and the software Zeiss Zen 2.3. Images from the first four sampling time points were taken using an exposure of 400 ms, while images of samples at 4 and 24 hpi were taken using an exposure of 350 ms and/or 400 ms, respectively. From each independent culture and time point at least five images, containing a total of more than 200 individual cells, were taken for image analysis.

Images were processed using ImageJ/FIJI (30). Values were normalized against the exposure time. A mask was created using the IsoData threshold method on the phase contrast channel and applied to the fluorescence channel, when needed the mask was adjusted to eliminate any frameshift. The binary process “Open” was applied. Particles were analyzed with the size parameter 0.80 - 2.00 µm^2^, and excluding particles touching the border. The average fluorescence intensity of each cell was determined and corrected by subtracting the average background intensity of the respective image. The background intensity of each image was assessed by inverting the previous mask and measuring the median intensity of the image. An average of 678 cells were measured for each sample, with the lowest number of measured cells being attributed to the second biological replicate at 0 hpi (205 cells).

### Data analysis

Data analysis and visualisation was performed using the package *tidyverse* from the *R* software (31, 32). The area under the curve (AUC) of the optical density and fluorescence was determined using the function “auc” from the integrated *R* package *MESS*. Fitness effects in competition experiments were determined by the ratio between AUC under a certain condition compared to the mean AUC of a strain growing as monoculture. Where appropriate, data were fitted into robust simple linear regression models using the function “lmrob” from the *R* package *robustbase* (33). Pearson’s correlations were determined using the function “cor” from the *R* package *stats*. Generalised linear mixed models (GLMM) were performed with the *R* package *gamlss* (34). One- and two-way ANOVAs were performed using the function “aov” from the *R* package *stats. Post-hoc* group comparisons were made with the *R* package *emmeans* (35).

## Acknowledgments

This work was supported by Marsden Fast Start grant 17-UOC-057 to MRE. RS was supported by a New Zealand International Doctoral Scholarship and a University of Canterbury PhD scholarship.

## References

1. Baba T, Ara T, Hasegawa M, Takai Y, Okumura Y, Baba M, Datsenko KA, Tomita M, Wanner BL, Mori H. 2006. Construction of Escherichia coli K-12 in-frame, single-gene knockout mutants: the Keio collection. Mol Syst Biol 2:2006.0008.

2. McFarland J. 1907. The nephelometer: An instrument for estimating the number of bacteria in suspensions used for calculating the opsonic index and for vaccines. JAMA XLIX:1176–1178.

3. Dalgaard P, Ross T, Kamperman L, Neumeyer K, McMeekin TA. 1994. Estimation of bacterial growth rates from turbidimetric and viable count data. Int J Food Microbiol 23:391–404.

4. Stevenson K, McVey AF, Clark IBN, Swain PS, Pilizota T. 2016. General calibration of microbial growth in microplate readers. Sci Rep 6:38828.

5. Guo R, McGoverin C, Swift S, Vanholsbeeck F. 2017. A rapid and low-cost estimation of bacteria counts in solution using fluorescence spectroscopy. Anal Bioanal Chem 409:3959–3967.

6. Chalfie M, Tu Y, Euskirchen G, Ward WW, Prasher DC. 1994. Green fluorescent protein as a marker for gene expression. Science 263:802–805.

7. Bloemberg GV, O’Toole GA, Lugtenberg BJ, Kolter R. 1997. Green fluorescent protein as a marker for Pseudomonas spp. Appl Environ Microbiol 63:4543–4551.

8. Shaner NC, Campbell RE, Steinbach PA, Giepmans BNG, Palmer AE, Tsien RY. 2004. Improved monomeric red, orange and yellow fluorescent proteins derived from Discosoma sp. red fluorescent protein. Nat Biotechnol 22:1567–1572.

9. Goedhart J, von Stetten D, Noirclerc-Savoye M, Lelimousin M, Joosen L, Hink MA, van Weeren L, Gadella TWJ Jr, Royant A. 2012. Structure-guided evolution of cyan fluorescent proteins towards a quantum yield of 93%. Nat Commun 3:751.

10. Lambert TJ. 2019. FPbase: a community-editable fluorescent protein database. Nat Methods 16:277–278.

11. Shaner NC, Steinbach PA, Tsien RY. 2005. A guide to choosing fluorescent proteins. Nat Methods 2:905–909.

12. Shaner NC, Patterson GH, Davidson MW. 2007. Advances in fluorescent protein technology. J Cell Sci 120:4247–4260.

13. Wilson E, Okuom M, Kyes N, Mayfield D, Wilson C, Sabatka D, Sandoval J, Foote JR, Kangas MJ, Holmes AE, Sutlief AL. 2018. Using fluorescence intensity of enhanced green fluorescent protein to quantify Pseudomonas aeruginosa. Chemosensors (Basel) 6(2): 21.

14. Schlechter RO, Jun H, Bernach M, Oso S, Boyd E, Muñoz-Lintz DA, Dobson RCJ, Remus DM, Remus-Emsermann MNP. 2018. Chromatic Bacteria - A broad host-range plasmid and chromosomal insertion toolbox for fluorescent protein expression in bacteria. Front Microbiol 9:3052.

15. Leveau JH, Lindow SE. 2001. Predictive and interpretive simulation of green fluorescent protein expression in reporter bacteria. J Bacteriol 183:6752–6762.

16. Tecon R, Binggeli O, van der Meer JR. 2009. Double-tagged fluorescent bacterial bioreporter for the study of polycyclic aromatic hydrocarbon diffusion and bioavailability. Environ Microbiol 11:2271–2283.

17. Ahn H-J, La H-J, Forney LJ. 2006. System for determining the relative fitness of multiple bacterial populations without using selective markers. Appl Environ Microbiol 72:7383– 7385.

18. Ross-Gillespie A, Gardner A, West SA, Griffin AS. 2007. Frequency dependence and cooperation: theory and a test with bacteria. Am Nat 170:331–342.

19. Friedman J, Higgins LM, Gore J. 2017. Community structure follows simple assembly rules in microbial microcosms. Nat Ecol Evol 1:109.

20. Remus-Emsermann MNP, Schmid M, Gekenidis M-T, Pelludat C, Frey JE, Ahrens CH, Drissner D. 2016. Complete genome sequence of Pseudomonas citronellolis P3B5, a candidate for microbial phyllo-remediation of hydrocarbon-contaminated sites. Stand Genomic Sci 11:75.

21. Choi K-H, Schweizer HP. 2006. mini-Tn7 insertion in bacteria with single attTn7 sites: example Pseudomonas aeruginosa. Nat Protoc 1:153–161.

22. Schada von Borzyskowski L, Remus-Emsermann M, Weishaupt R, Vorholt JA, Erb TJ. 2015. A set of versatile brick vectors and promoters for the assembly, expression, and integration of synthetic operons in Methylobacterium extorquens AM1 and other alphaproteobacteria. ACS Synth Biol 4:430–443.

23. Schlechter R, Remus-Emsermann M. 2019. Delivering “Chromatic Bacteria” Fluorescent Protein Tags to Proteobacteria Using Conjugation. BIO-PROTOCOL 9.

24. Kremers G-J, Goedhart J, van Munster EB, Gadella TWJ Jr. 2006. Cyan and yellow super fluorescent proteins with improved brightness, protein folding, and FRET Förster radius. Biochemistry 45:6570–6580.

25. Bindels DS, Haarbosch L, van Weeren L, Postma M, Wiese KE, Mastop M, Aumonier S, Gotthard G, Royant A, Hink MA, Gadella TWJ Jr. 2017. mScarlet: a bright monomeric red fluorescent protein for cellular imaging. Nat Methods 14:53–56.

26. Kolek J, Sedlar K, Provaznik I, Patakova P. 2016. Dam and Dcm methylations prevent gene transfer into Clostridium pasteurianum NRRL B-598: development of methods for electrotransformation, conjugation, and sonoporation. Biotechnol Biofuels 9:14.

27. Tancos MA, Chalupowicz L, Barash I, Manulis-Sasson S, Smart CD. 2013. Tomato fruit and seed colonization by Clavibacter michiganensis subsp. michiganensis through external and internal routes. Appl Environ Microbiol 79:6948–6957.

28. Cormack BP, Valdivia RH, Falkow S. 1996. FACS-optimized mutants of the green fluorescent protein (GFP). Gene 173:33–38.

29. Harder W, Attwood MM, Quayle JR. 1973. Methanol Assimilation by Hyphomicrobium sp. Journal of General Microbiology 78:155–163.

30. Schindelin J, Arganda-Carreras I, Frise E, Kaynig V, Longair M, Pietzsch T, Preibisch S, Rueden C, Saalfeld S, Schmid B, Tinevez J-Y, White DJ, Hartenstein V, Eliceiri K, Tomancak P, Cardona A. 2012. Fiji: an open-source platform for biological-image analysis. Nat Methods 9:676–682.

31. R Core Team, Others. 2013. R: A language and environment for statistical computing. Vienna, Austria. URL https://www.R-project.org/.

32. Martin G. 2021. Introduction to the tidyverse. An Introduction to Programming with R. Journal of Open Source Software 4(43):1686. URL https://doi.org/10.21105/joss.01686.

33. Rousseeuw P, Croux C, Todorov V, Ruckstuhl A, Salibian-Barrera M, Verbeke T, Koller M, Maechler M. 2012. robustbase: Basic Robust Statistics. R package version 0.9-7. URL http://robustbase.r-forge.r-project.org.

34. Rigby RA, Stasinopoulos DM. 2005. Generalized additive models for location, scale and shape (with discussion). Journal of the Royal Statistical Society: Series C (Applied Lenth RV. 2020. emmeans: Estimated Marginal Means, aka Least-Squares Means. R package version 1.5.3. URL https://github.com/rvlenth/emmeans.

35. Remus-Emsermann MNP, Kim EB, Marco ML, Tecon R, Leveau JHJ. 2013. Draft genome sequence of the phyllosphere model bacterium Pantoea agglomerans 299R. Genome Announc 1(1): e00036–13.

36. Innerebner G, Knief C, Vorholt JA. 2011. Protection of Arabidopsis thaliana against leaf-pathogenic Pseudomonas syringae by Sphingomonas strains in a controlled model system. Appl Environ Microbiol 77:3202–3210.

