## Supplemental material for "Fluorescent protein expression as a proxy of bacterial fitness in a high throughput assay"

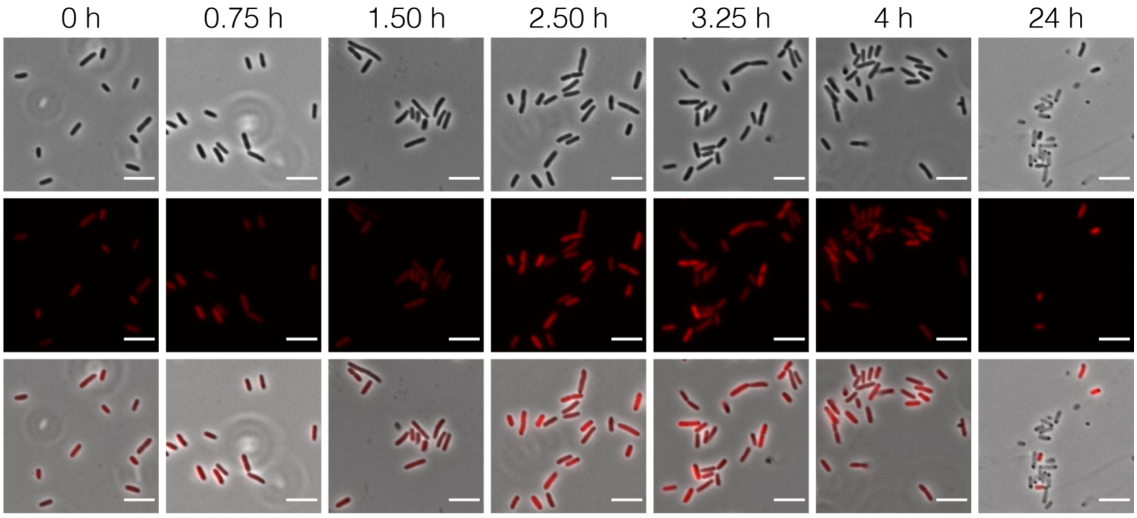


**Figure S1. Pe::red single-cell fluorescence over time.** Representative images of Pe::red cultures grown in nutrient broth. Samples were taken at 0, 0.75, 1.50, 2.50, 3.25, 4, and 24 h post-inoculation. **Top row**: phase contrast; **middle row**: red channel; **bottom row**: overlay between phase contrast and red channel images. Scale bar: 5 µm.

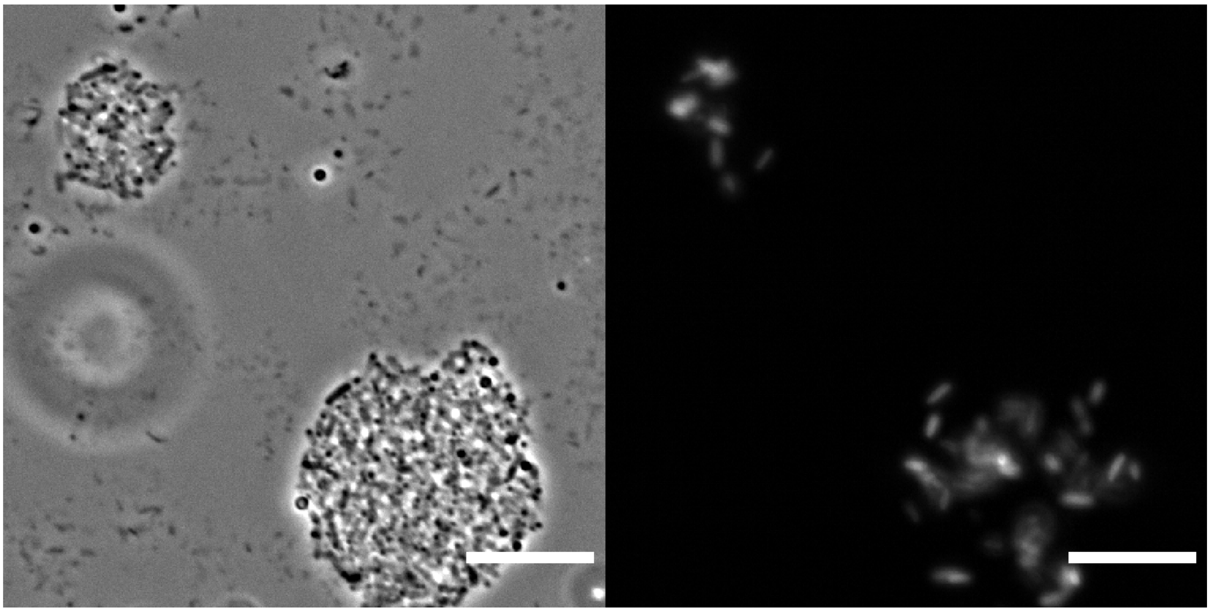

**Figure S2.** Representative image of Pe::red cell aggregates. Sample of NB-grown Pe::red taken at 24 h post-inoculation in conical flasks. **(Left panel)** Phase contrast image. **(Right panel)** Red fluorescence image. Scale bars 10 µm.

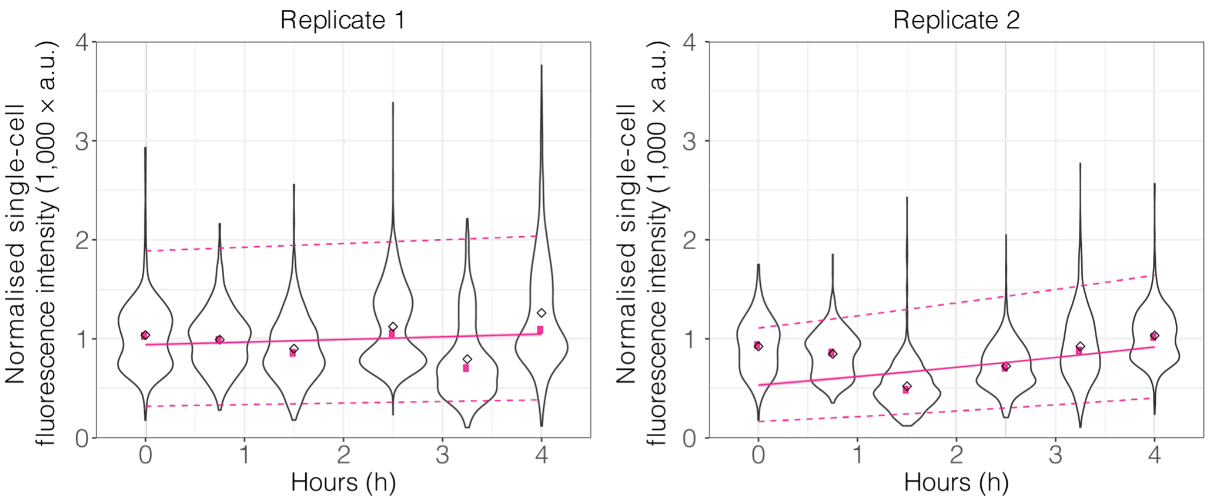


**Figure S3. Single cell fluorescence in growing Pe::red cultures.** Two independent cultures of NB-grown Pe::red in conical flasks were sampled at 0, 0.75, 1.50, 2.50, 3.25 and 4 h post-inoculation. Fluorescence intensity was measured from individual cells. Data was fitted to a log-normal distribution. . Replicate 1, $p$-value < 0.05, adjusted $R^2$ = 0.0079; Replicate 2, $p$-value < 0.05, adjusted $R^2$ = 0.13. Empty diamond: mean of each group; filled coloured circle: median of each group; continuous line: regression line; dashed lines: prediction interval (upper and lower limit).

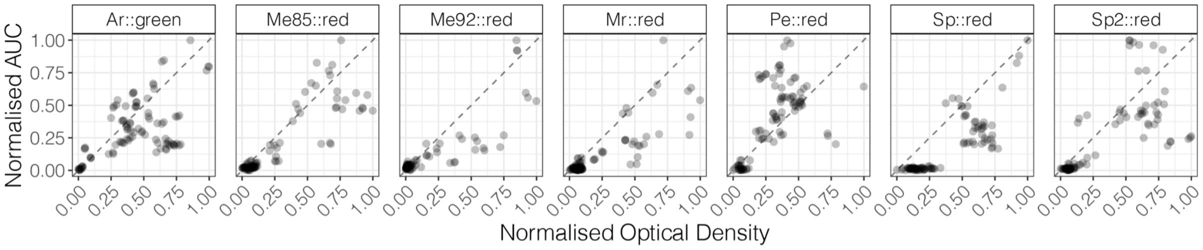


**Figure S4. Correlation between optical density and fluorescence in different strains and different growth media.** Each point represents a biological replicate (individual strain grown in a defined media). Min-max normalisation was applied to optical density measurements and AUC from fluorescent data, where 0 and 1 represent the minimal and maximal value for each dataset. Dashed line represents the identity line (y = x). Pearson's correlation = 0.72. GLMM, $p$-value < 0.05, *pseudo*-$R^2$ = 0.80.

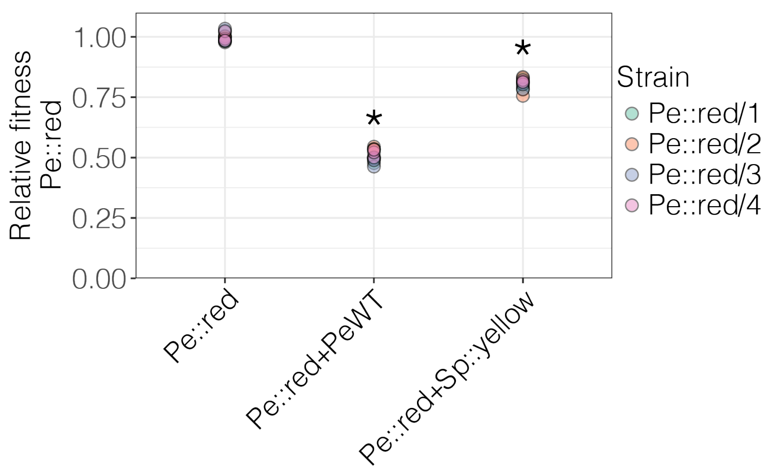


**Figure S5. Fitness of independent Pe::red strains.** Fitness of four independent Pe::red Tn*7* insertion mutants in competition with a PeWT and Sp::yellow. Monocultures (Pe::red) were included as control and to calculate the relative fitness of each strain based on area under the fluorescence curves (AUC). Two-way ANOVA was used to compare fitness differences between strains and treatments (monocultures and growth in the presence of a competitor, PeWT or Sp::yellow), $p\text{-value}_{\text{Strain}} = 0.256$; $\text{p-value}_{\text{Treatment}} < 0.05$; $\text{p-value}_{\text{Interaction}} = 0.238$. Asterisk (*) indicates statistical significance from a *post-hoc* Bonferroni test (alpha = .95, *p*-value < 0.05) relative to the monoculture treatment.

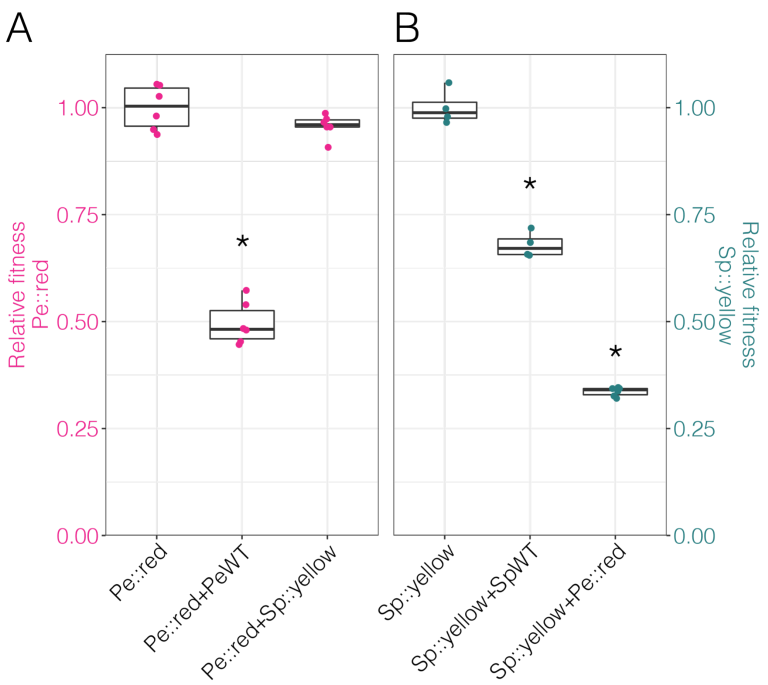


**Figure S6. Competition in minimal medium.** The fluorescently-tagged bacterial strains Pe::red and Sp::yellow were co-inoculated in minimal medium supplemented with 0.2% w/v succinate and their growth was estimated using fluorescence intensity. The data was background subtracted to normalise against the autofluorescence of the media. **(A)** Relative fitness of Pe::red in monoculture, co-culture with PeWT, and co-culture with Sp::yellow. Fitness determined as the AUC derived from red fluorescence relative to Pe::red in monoculture. **(B)** Relative fitness of Sp::yellow in monoculture, co-culture with SpWT, and co-culture with Pe::red. Relative fitness of Sp::yellow was determined as the AUC derived from yellow fluorescence relative to Sp::yellow in monoculture. Groups were compared using one-way ANOVA test, where * = $p$-value < 0.05 compared to the respective monoculture (Pe::red or Sp::yellow in **(A)** or **(B)**, respectively) using a Bonferroni *post-hoc* test.
